# Investigating BPDE-induced embryonic toxicity employing hiPSC-based models

**DOI:** 10.1101/2024.12.16.628678

**Authors:** Vanessa Cristina Meira de Amorim, Leon Szepanowski, Alessia Lofrano, Christiane Loerch, Wasco Wruck, Nina Graffmann, James Adjaye

**Affiliations:** Institute for Stem Cell Research and Regenerative Medicine, Medical Faculty and University Hospital Düsseldorf, Heinrich-Heine University Düsseldorf, Düsseldorf, 40225, Germany; Department of Ophthalmology, University Hospital Duesseldorf, Duesseldorf, Germany; University College London (UCL) –EGA Institute for Women’s Health, Zayed Centre for Research into Rare Diseases in Children (ZCR). 20 Guilford Street, London WC1N 1DZ

**Keywords:** Nijmegen Breakage Syndrome (NBS), benzo[a]pyrene diol epoxide (BPDE), developmental toxicology, DNA damage response, induced pluripotent stem cells, neuroprogenitor cells

## Abstract

Benzo[a]pyrene diol epoxide (BPDE) is a metabolite of the environmental contaminant Benzo[a]pyrene- a byproduct of incomplete combustion of organic matter. BPDE reacts with DNA to form BPDE-DNA bulky adducts which if not removed can lead to mutations due to DNA base-pair substitutions. While the effects of BPDE on somatic cells are fairly well described, its effects on early human development are currently unknown. In this study, we investigated for the first time the effect of BPDE on human induced pluripotent stem cells (hiPSCs) and their differentiated neuroprogenitor cells (NPCs) as a model for early embryonic development. Furthermore, we compared hiPSCs and NPCs derived from cells of patients suffering from Nijmegen Breakage Syndrome (NBS), which is a chromosomal instability disorder characterized by defective DNA repair and increased risk of malignancies. Transcriptome analysis, coupled with protein content analysis employing immunostaining and Western blots, revealed that hiPSCs are more sensitive to BPDE exposure when compared to NPCs with an enhanced expression of several genes associated with p53-mediated DNA damage response, including DNA repair by lesion bypass, cell cycle checkpoints and extrinsic apoptosis. We also identified that cells from NBS patients showed less apoptotic response and a distinct p53 response than their healthy counterparts. This iPSC-based study enhances our meagre knowledge of the effects of BPDE on early human development in both healthy individuals and NBS patients. Furthermore, our model conforms with the 3Rs principle.

## 1. Introduction

There is currently a worldwide endeavor to minimize and even replace animal experimentation. There are numerous problems associated with animal experimentation which include ethical issues, the high costs of animal experiments and difficulties in recapitulating specific human pathologies. The pharmaceutical and chemical industries contend with a failure rate of up to 90% in translating animal effects to human subjects (1,2).

Developmental toxicity testing aims to identify substances that can cause disturbances during embryo-fetal development. Currently, the gold standard are animal-based assays which are time consuming, expensive and do not always reliably translate to human physiology (3). Besides developmental toxicological evaluation, the identification and investigation of the effects of DNA-damaging agents (genotoxins) is of high importance. While DNA-repair mechanisms remain mostly conserved among mammals, there are marked differences on the efficiency of such repair processes between distinct species (4). Tests *in vitro* with a high sensitivity are also part of the standard battery of tests used for genotoxic assessment in human pharmaceuticals (5), but their specificity is often unsatisfactory (6,7).

A promising alternative for animal experimentation is the use of human pluripotent stem cells (hiPSCs) and their differentiated progeny as *in vitro* substitutes. In 2023, the U.S. Congress approved the FDA Modernization Act 2.0, stating that drug developers can now propose alternative methods, including the use of hiPSC-derived models, for the assessment of drug safety during the pre-clinical phase instead of mandating the use of animal models (8). hiPSC-based models are particularly interesting alternatives for developmental toxicology testing since they can mimic the developing embryo and parts of organ development (9).

Furthermore, hiPSCs are advantageous for modeling specific diseases, particularly genetic disorders. One example is Nijmegen Breakage Syndrome (NBS), a chromosomal instability disorder, characterized by an impaired immune system, microcephaly, growth retardation, premature aging, and increased susceptibility to malignancies (10). It is caused by a mutation within the NBS gene that codes for the protein Nibrin (NBN), which is part of the MRN complex together with Meiotic recombination 11 (MRE11) and RAD50 Double Strand Break Repair Protein (RAD50). This complex is involved in DNA damage signaling and repair, particularly of DNA double-strand breaks, a pathway that is severely impaired in NBS patients. Due to this deficiency in DNA repair, NBS patients have marked chromosomal instability, developing breakages and chromosomal rearrangements (11). That in turn leads to the development of malignancies, with over 40% of NBS patients developing one by the age of 20, predominantly hematological cancers (10).

Benzo[a]pyrene (B[a]p) is a polycyclic aromatic hydrocarbon and environmental contaminant formed during incomplete combustion of organic matter, such as exhaust fumes and cigarette smoke. B[a]p is metabolized in the body into benzo[a]pyrene diol epoxide, or BPDE, a potent mutagen and carcinogen. BPDE forms bulky DNA adducts by binding to the N2 atom of guanine (dG-N2-BPDE adduct) which, if not removed by nucleotide excision repair (NER), can lead to base-pair substitution mutations via trans-lesion DNA synthesis (TLS) (12) resulting in tumorigenesis (13). B[a]p is lipophilic and can cross the placenta to reach the developing fetus. The placenta can metabolize B[a]p into BPDE, and the fetuses themselves are metabolically capable, which can lead to fetal genotoxic exposure (14–16). Furthermore, BPDE-DNA adducts are detectable in the sperm (17) and ovarian cells (18) of cigarette smokers, and these DNA modifications can be paternally transmitted through the spermatozoa to the embryo (19). The potential issues that could arise from the presence of BPDE adducts on pre-implantation embryos and fetuses, are currently unknown. The undifferentiated pluripotent stem cells represent the pre-gastrulation stage whilst the differentiated neural progenitor cell gastrulation towards ectoderm.

Our work is the first to perform a comparative analysis of the response to BPDE exposure in pluripotent stem cells and their differentiated neural progenitor cell (NPC) progeny, using both healthy cells and those with an NBS mutation. We show here for the first time that low doses of BPDE do not interfere with the maintenance of the pluripotent state. However, it has distinct impacts on p53-associated stress responses such as DNA-damage repair and apoptosis in iPSCs and NPCs.

## 2. Methods and materials

### 2.1. Stem Cell Cultivation

Two hiPSC lines from healthy individuals, UM51 and iPSC-12, and one hiPSC line from an individual suffering from NBS, NBS8, were used in this study (Supplementary Table 1). hiPSCs were cultivated on plates coated with Matrigel (Corning) and fed with mTeSR plus™ medium (StemCell Technologies). Cell passaging was done every 5-6 days with ReLeSR™ (Stem Cell Technologies) according to the manufacturer’s instructions. UM51 (20) and NBS8 (21) were generated under the ethical approval of the Ethikkommission der Medizinischen Fakultät der Heinrich-Heine-Universität Düsseldorf. iPSC-12 is a commercially available iPSC line from Cell Applications Inc. (https://www.cellapplications.com/).

#### Neural Progenitor Cell Differentiation

For NPC differentiation, hiPSCs were cultivated until 75-90% confluence. The cells were dissociated into single cells with accutase (Sigma-Aldrich) and plated in a 96-well U-bottom, low attachment plate (Thermo Scientific) with 100μL mTeSR Plus™ supplemented with 10 μM ROCK inhibitor (Y-27632, Sigma-Aldrich) in a density of 10^4^ cells/well. The cells were incubated at 37°C with 5% CO2 for 24 hours, after which 50 μl of the medium was aspirated and replaced by 100 μl of neural induction medium (NiM, consisting of 47% DMEM/F12 (Gibco), 47% Neurobasal Medium™ (Gibco), 2% B27™ w/o retinoic acid (Gibco), 1% N2 (Gibco), 1% GlutaMAX™ (Gibco), 1% MEM non-essential amino acids (NEEA) (Gibco) and 1% Penicillin/Streptomycin (Gibco). In the following 5 days, 100 μl of medium was replaced daily with fresh NIM supplemented with 10 μM SB-431542 (Tocris), 500 nM LDN-193189 (Merck) and, until day 3, 10 μM ROCK inhibitor. The resultant neurospheres were seeded in Growth Factor Reduced Matrigel® (Corning) coated 6-well plates and fed daily with Neural Differentiation Medium (NDM, consisting of 95% Neurobasal Medium™ (Gibco), 2% B27™ w/o retinoic acid (Gibco), 1% N2 (Gibco), 1% GlutaMAX™ (Gibco), and 1% Penicillin/Streptomycin (Gibco) supplemented with 20 ng/ml epidermal growth factor (EGF) and 20 ng/ml fibroblast growth factor 2 (FGF2). On day 18, neural rosettes were selected with STEMdiff™ Neural Rosette Selection Reagent (Stem Cell Technologies) as per the manufacturer’s recommendation. The rosettes were then incubated with accutase for 30 min at 37°C and the resulting cell aggregates were seeded at a density of 1:4 on Growth Factor Reduced Matrigel® (Corning) coated 6-well plates for expansion and fed daily with NDM supplemented with 20 ng/ml EGF and 20 ng/ml FGF2. For passaging, the NPCs were incubated with accutase for 5 min at 37°C and the resulting aggregates were seeded at a density of 1:4 on Growth Factor Reduced Matrigel® (Corning) coated 6-well plates every 5-6 days.

### 2.2. BPDE treatment

BPDE (Santa Cruz Biotechnology) was dissolved in DMSO (Sigma-Aldrich) at a concentration of 40mM and the aliquots were stored at −80°C for up to six months. For treatment of hiPSCs and NPCs cells were cultured until 60-70% confluent, and then medium replaced with fresh medium containing BPDE (25nM or 75nM) or vehicle control (0.02% DMSO) for 24h. Thereafter, medium was discarded, and cells processed for analysis.

### 2.3. Resazurin reduction assay

A 0.15 mg/ml stock solution of resazurin (Sigma-Aldrich) was prepared by dissolving 5mg of resazurin in 50mL of sterile PBS and kept at 4°C until use. Cells were cultivated in triplicates on a 96-well plate and continuously exposed to BPDE in concentrations ranging from 10nM to 21μM for 22h, when 10μL medium supplemented with resazurin stock solution was added to the culture in a 1:10 dilution. The plates were returned to 37°C and incubated for a further 2h. Living cells convert resazurin into resofurin, a fluorescent compound, and fluorescence was measured using a microplate fluorimeter (Eppendorf PlateReader AF2200) equipped with a filter set of 560nm excitation and 590nm emission. Measurements of cell cultures treated with increasing doses of BPDE and normalized according to the manufacturer’s protocol were imported into the R environment (22). The packages dr4pl (23) and ggplot2 (24) were employed to use a logistic model for curve-fitting, plotting the curve, and calculating the IC50 and additionally IC80 and IC90.

### 2.4. Immunocytochemistry

Cells were fixed with 4% paraformaldehyde (PFA) for 15 minutes. For staining of intracellular proteins, the cells were permeabilized with 0.5% Triton-x-100/DPBS for 10 min and then washed 2x with DPBS. Cells were then incubated with blocking buffer containing 3% bovine serum albumin (BSA, Sigma-Aldrich)/DPBS for 1h at RT. After the blocking period, cells were incubated with primary antibody diluted in blocking buffer overnight at 4°C with shaking. In the morning the cells were then washed 3x/5min with DPBS, then further incubated with a secondary antibody solution diluted in blocking buffer, supplemented with the nuclear stain Hoechst 33258 (Thermo Fisher) for 2h RT (antibodies are listed in Supplementary Table 2. Pictures were taken using a Zeiss LSM 700 microscope and analyzed using the software Zen Blue 2.5. When applicable, cells were counted manually using the program image J (ver. 1.54) and ratios of treated conditions and control were calculated using Microsoft Excel, as were standard deviations (SD). Ratios and SD were visualized as bar graphs. Statistical significance was measured using 2-way ANOVA and Tukey’s multiple comparison test using GraphPad Prism 8. P values ≤ 0.05 were considered statistically significant.

### 2.5. Propidium Iodide (PI) staining and cell cycle FACS analysis

hiPSCs were treated with BPDE for 24h. Thereafter, cells were harvested using accutase (Sigma-Aldritch) and centrifuged at 500xg for 5min at 4°C. 10^5^ cells were transferred to FACS tubes (Corning), washed with 1ml of PBS and centrifuged as stated above. The supernatant was discarded, and the cells were incubated for 1h in 25μl of staining solution composed of 0.1% sodium citrate, 0.1% Triton-x-100 and 50mg/L of propidium iodide (PI) (Invitrogen), diluted in distilled water. Cell cycle measurements were carried out on a CytoFlex BA26183 from Beckman Coulter and analysed with CytExpert 2.3. Statistical significance was measured using 2-way ANOVA and Tukey’s multiple comparison test using GraphPad Prism 8. P values ≤ 0.05 were considered statistically significant.

### 2.6. Reverse Transcripion and real time PCR

TRIzol® was used to extract RNA from treated and non-treated hiPSCs and NPCs. Total RNA was extracted using the Direct-zol^TM^ RNA MiniPrep kit (Zymo Research) according to the manufacturer’s instructions. To avoid DNA contamination, a 30 min treatment with DNase was applied. The isolated RNA was transcribed into cDNA using the Reverse Transcription TaqMan® Kit (Applied Biosystems) following the manufacturer’s instructions. The real time PCR was performed with Power SYBR® Green (Applied biosystems) on a ViiA7 machine (Applied biosystems). Mean values were normalized to the housekeeping gene *RPLP0* and the cycle threshold (CT) for each sample was determined with the ViiA7 Software v1.2 from Applied Biosystems. Fold-change values are depicted as mean values with 95% confidence interval. Statistical significance was measured using 2-way ANOVA and Tukey’s multiple comparison test using GraphPad Prism 8. P values ≤ 0.05 were considered statistically significant. Primer sequences used are listed in Supplementary Table 3.

### 2.7. 5-ethynyl-2’-deoxyuridine (EdU) incorporation assay

hiPSCs were treated for 24h with BPDE. EdU incorporation and the fluorescence staining for incorporated EdU was performed using the Click-iT™ EdU Cell Proliferation Kit for Imaging, Alexa Fluor™ 488 dye (Invitrogen), as instructed by the manufacturer. Hoechst 33258 was used for DNA staining. Pictures were taken using a Zeiss LMS 700 and analyzed using the software Zen Blue 2.5. EdU+ cells were counted manually using the program image J (ver. 1.54) and ratios of treated conditions and control were calculated using Microsoft Excel 2010, as were standard deviations (SD). Ratios and SD were visualized as bar graphs. Statistical significance was measured using 2*way ANOVA and Turkey’s multiple comparison test using GraphPad Prism 8. P values ≤ 0.05 were considered statistically significant.

### 2.8. Analysis of Gene Expression Data

Total RNA was extracted using the Direct-zol^TM^ RNA MiniPrep kit (Zymo Research) according to the manufacturer’s instructions from iPSCs UM51, iPSC-12 and NBS8, in CTRL conditions and after 24h treatment with 75nM BPDE. RNA was sent for analysis at the Biomedizinisches Forschungszentrum (BMFZ) facility at Heinrich-Heine University, Duesseldorf. Affymetrix CEL files were imported into the R/Bioconductor (22) environment,background-corrected, and normalized using the Robust Multi-array Average (RMA) method from the package oligo (25). Genes were considered expressed when their detection p-values - calculated as described in Graffmann et al. (26) - were below a threshold of 0.05. Using these expressed genes, expression was dissected with Venn diagrams employing the R package “VennDiagram” (27). Tables of Pearson correlation coefficients were generated using the R-built-in method “cor”. Hierarchical clustering dendrograms were generated with the method “hclust” using Pearson correlation as similarity measure and “complete linkage” as cluster agglomeration method. The “heatmap.2” function from the “gplots” package (28) was applied either with Pearson correlation as similarity measure and color scaling per gene or with Euclidean distance as distance measure and color scaling over the whole heatmap. Genes from the intersection of the venn diagrams, i.e., expressed in both conditions were further filtered for up-regulation by a ratio > 1.5 and down-regulation by a ratio < 0.67. When there were replicates, a threshold of 0.05 for the p-value of the differential expression test from the Bioconductor “limma” package (29) was added to the filter criteria.

### 2.9. Analysis of Pathways and Gene Ontologies (GOs)

Differentially up- and down-regulated genes with fold changes of 1.5 or 0.6 and P-value 0,05 were subjected to GO analysis via the Bioconductor package “GOstats” (**30**). KEGG pathways and genes associated with them were downloaded from the KEGG database (**31**) and over-represented KEGG pathways were calculated for up- and down-regulated genes employing the R-built-in hypergeometric test. The enrichment analysis tool Metascape (**32**) was used to compare input gene lists extracted from the microarray data to thousands of available gene sets defined by their involvement in biological processes, protein localization, enzymatic function, pathways among other features.

### 2.10. Western Blotting

RIPA buffer (Sigma-Aldrich), with added protease and phosphatase inhibitors (Roche,), was used to extract the total protein from hiPSCs and NPCs. Protein concentration was measured with the Pierce™ BCA Protein Assay Kit (Thermo Fisher). 20µg of protein was run through a 4%-12% SDS-PAGE gel (NuPAGE, Thermo Fisher) and transferred to a 0.45 µm nitrocellulose membrane (GE Healthcare) using wet blotting. The membrane was blocked with 5% milk in TBST (20mM Tris, 150nM NaCl and 0,05% Tween 20) for 1h and stained for primary antibody in 4°C overnight, under constant agitation. The membranes were subsequently washed three times with TBST and stained for secondary antibody for two hours RT. Details of the antibodies used can be seen in Supplementary Table 2. Anti ß-actin and anti-RPLP0 were used as housekeepers. Protein bands were detected with Pierce™ ECL Western Blotting Substrate. Signaling was visualized using FusionCapt Advance FX7 and band intensities were quantified in the software Fusion Capt Advance (PeqLab) using rolling ball background correction.

## 3. Results

### 3.1. 24h of BPDE exposure does not impact hiPSC pluripotency

In this study, we aimed at elucidating the effects of the genotoxin BPDE on iPSCs and see if iPSCs derived from an NBS patient (NBS8) react differently towards BPDE- induced DNA damage than wild type (WT) control cells (UM51, iPSC-12). To test for BPDE effects on the cellular stress response, we selected a suitable BPDE dosage scheme that assures cellular viability of more than 80% and does not interfere with pluripotency. Therefore, we treated all three hiPSC lines for 24h with distinct doses of BPDE and measured viability via resazurin assay. From this, we selected 25nM and 75nM as the IC10 and IC20, respectively, which were used for all further experiments. (Supplementary figure 1).

The hiPSC lines UM51, iPSC-12, and NBS8 were exposed to 25 or 75nM BPDE for 24h or kept in control conditions, then subjected to microarray analysis. To investigate whether BPDE exposure had an effect on pluripotency, microarray data was used to generate a heatmap of twelve genes which are key in the regulation of pluripotency (Figure 1A). Genes such as POU Class 5 Homeobox 1 (*POU5F1*), SRY-Box Transcription Factor 2 (*SOX2*) and *NANOG,* key players orchestrating the maintenance of pluripotency, remained highly expressed after treatment, while genes involved in hiPSC differentiation like Gremlin 1 (*GREM1*) and bone morphogenetic protein 4 (*BMP4*) had a low expression. qRT-PCR for *POU5F1* (*OCT4*), *SOX2* and *NANOG* confirmed the microarray data (Figure 1B) and immunocytochemistry for OCT4 also revealed no difference in expression after BPDE treatment (Figure 1C).

**Figure 1:**
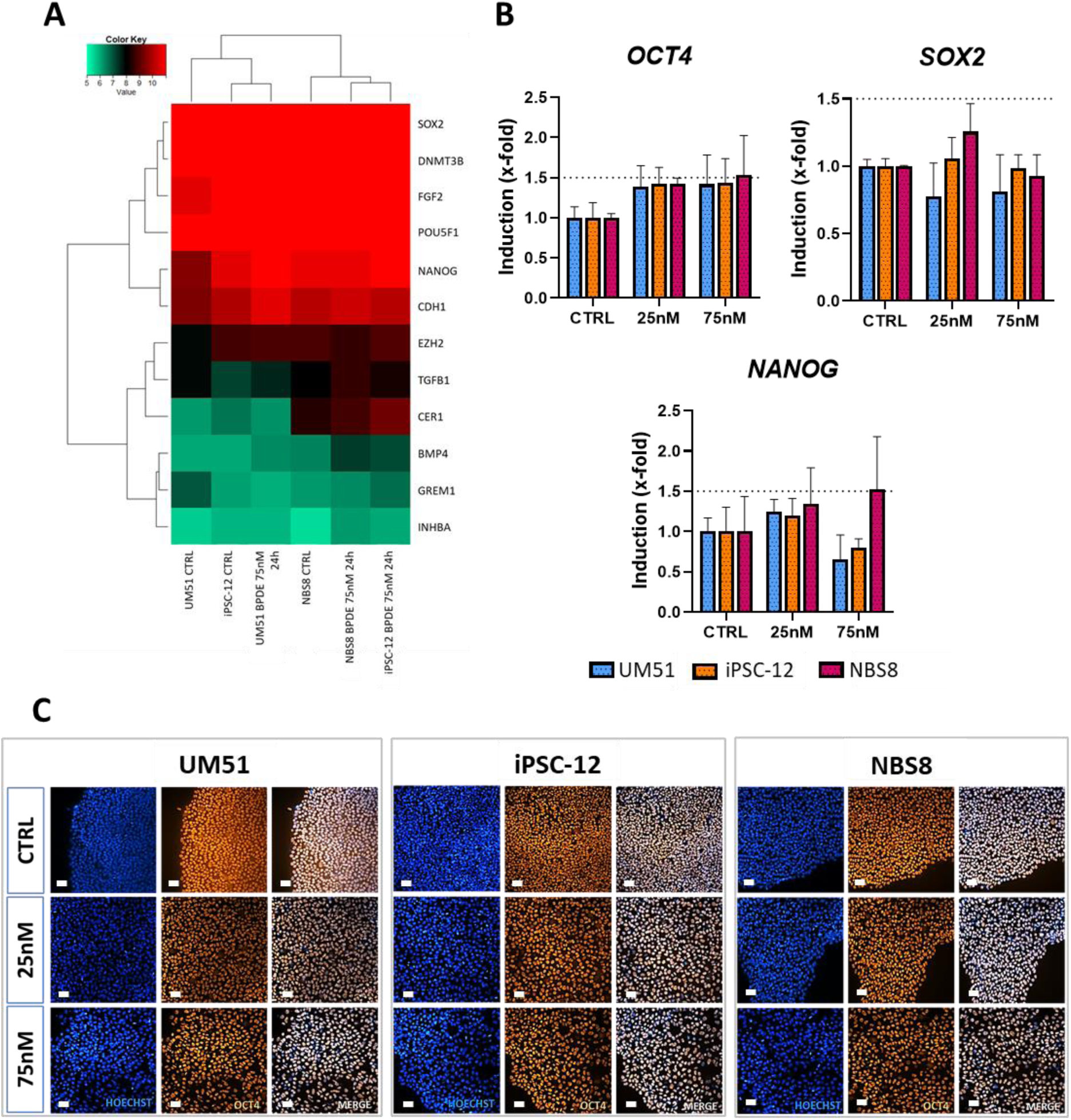
24h of BPDE exposure does not negatively impact pluripotency. WT and NBS8 hiPSCs were treated with 25nM and 75nM BPDE for 24h. RNA was extracted for microarray analysis or cells were fixed for immunocytochemistry. **(A)** Euclidean heatmap made with data extracted from the microarray analysis represents relative expression of key genes associated with pluripotency maintenance after 24h of 75nM BPDE exposure. **(B)** BPDE treatment did not alter *OCT4*, *SOX2* and *NANOG* transcription as confirmed via qRT-PCR (N=3. Error bar depicts 95% confidence interval. Scale bar 50μm. Dashed lines mark 1.5-fold.) **(C)** nor did it alter OCT4 expression as seen on immunocytochemistry.

### 3.2. BPDE exposure enhances gene expression of cell cycle arrest related genes, but does not impact cell cycle

As the integrity of the whole organism depends on the ability of stem cells to safely replace lost cells with their progeny, it is mandatory that stem cells keep their genome intact in order to avoid growth of malignantly transformed cells (33). Thus, they need to protect their genome by enhanced repair activity and if this is not possible, they usually die through apoptosis (34). To see if these pathways are differentially regulated in healthy and diseased cells, we first investigated if the cell cycle was affected by BPDE as cell cycle arrest is necessary to give the cells enough time for DNA repair.

To investigate the transcriptional changes induced by BPDE exposure regardless of the mutation, we pooled the microarray results of all samples into control and BPDE treated (Supplementary figure 2A). We also investigated the transcriptional downstream effects of the mutation, both in the absence of stimulus and after BPDE treatment, comparing the WT and NBS8 lines in control conditions (Supplementary figure 2B) and after genotoxic exposure (Supplementary figure 2C). We performed Gene Ontologies (GO), KEGG pathway, as well as Metascape analyses (Supplementary figures 3 to 6).

Gene ontology analysis of 513 genes exclusively expressed after BPDE treatment (Figure 2A) revealed, amongst others, the cluster “DNA damage response, signal transduction by p53 class mediator resulting in cell cycle arrest” with four regulated genes: cyclin dependent kinase inhibitor 1A (*CDKN1A*); Mucin1 (*MUC1*); polo like kinase 2 (*PLK2*) and proline rich acidic protein 1 (*PRAP1*). qRT-PCR for *CDKN1A* and Growth arrest and DNA damage inducible alpha (*GADD45A*), another p53-inducible DNA damage and cell cycle arrest associated gene, revealed that all three cell lines exhibited upregulated expression levels (mean > 3-fold) of *GADD45A* after exposure to 75nM BPDE, while there was no regulation after treatment with 25nM BPDE. *CDKN1A* was upregulated intensely in NBS8 (mean 1.6-fold at 25nM and 2.9- fold at 75nM) and slightly in iPSC-12 (mean 1.6-fold at 75nM) (Figure 2B). However, a cell cycle analysis using propidium iodide staining and flow cytometry showed that cell cycle distribution on all three lines was unaffected after BPDE exposure (Figure 2C). This result was supported by EdU staining, which confirmed that BPDE exposure had no effect on the percentage of cells in S phase (Figure 2D).

**Figure 2:**
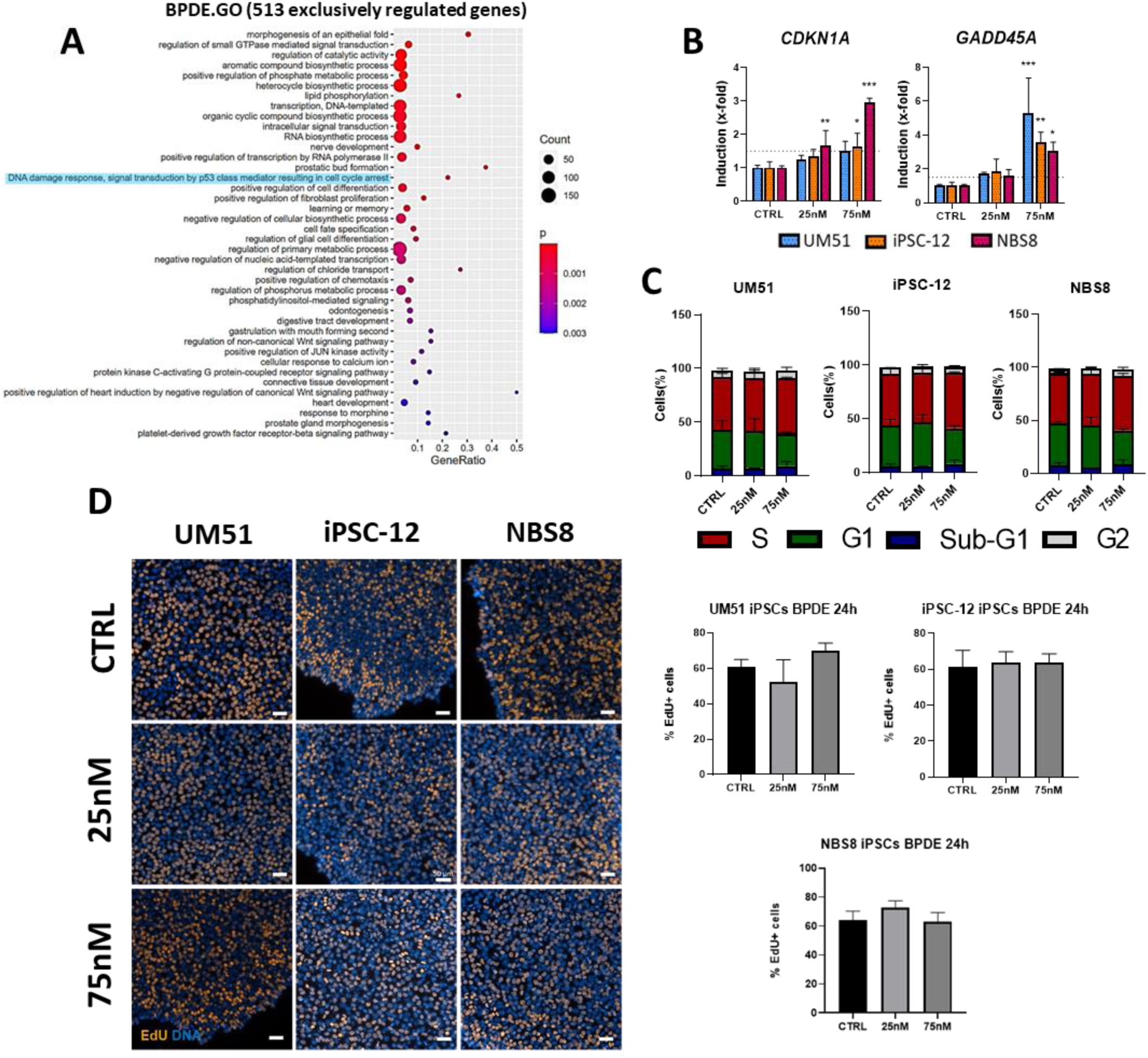
BPDE exposure enhances gene expression of cell cycle arrest related genes, but does not influence cell cycle. hiPSCs were treated for 24h with 25 or 75 nM BPDE **(A)** GO-Analysis for 513 genes exclusively expressed in all 3 cell lines after BPDE exposure in hiPSCs. The relevant DNA-damage cluster is highlighted in blue. **(B)** qRT-PCR for *CDKN1A* and *GADD45A.* Error bar depicts 95% confidence interval. N=3, *p<0.05, ** = p<0.01, *** = p<0.001. Dashed lines mark 1.5-fold. **(C)** Cell cycle distribution was analyzed via propidium iodide staining measured by FACS and presented here in percentage as bar graphs. N=3, mean +/- standard deviation. **(D)** Representative images of EdU staining of hiPSCs after CTRL, 25nM and 75nM treatment and bar graphs representing the percentage of EdU positive cells on each cell line compared to Hoechst after 24h treatment. N=3, mean +/- standard deviation. Scale bar 50μm.

### 3.3. BPDE treatment enhances expression of targets related to the extrinsic apoptotic pathway in hiPSCs

As we observed that the expression of genes related to cell cycle arrest was enhanced by BPDE while there was no actual cell cycle arrest detectable, we aimed to investigate if BPDE influences apoptosis in hiPSCs.

The GOs extracted from the 254 genes upregulated after BPDE treatment revealed several clusters related to both positive and negative regulation of cell death (Figure 3A). Notably, several genes related to the extrinsic apoptotic signaling pathway were upregulated including all four known receptors for TNF-related apoptosis-inducing ligand (TRAIL): TNF Receptor Superfamily Member 10A (*TNFRSF10A*), *TNFRSF10B*, *TNFRSF10C* and *TNFRSF10D.* qRT-PCR for *TNFRSF10A* (Figure 3B) confirmed its upregulation in all three lines (mean > 1.7- fold), while the expression of the intrinsic apoptosis genes BCL2 associated X, apoptosis regulator (*BAX*) and Bcl-2-binding component 3 (*BBC3*) remained unchanged (Figure 3B).

**Figure 3:**
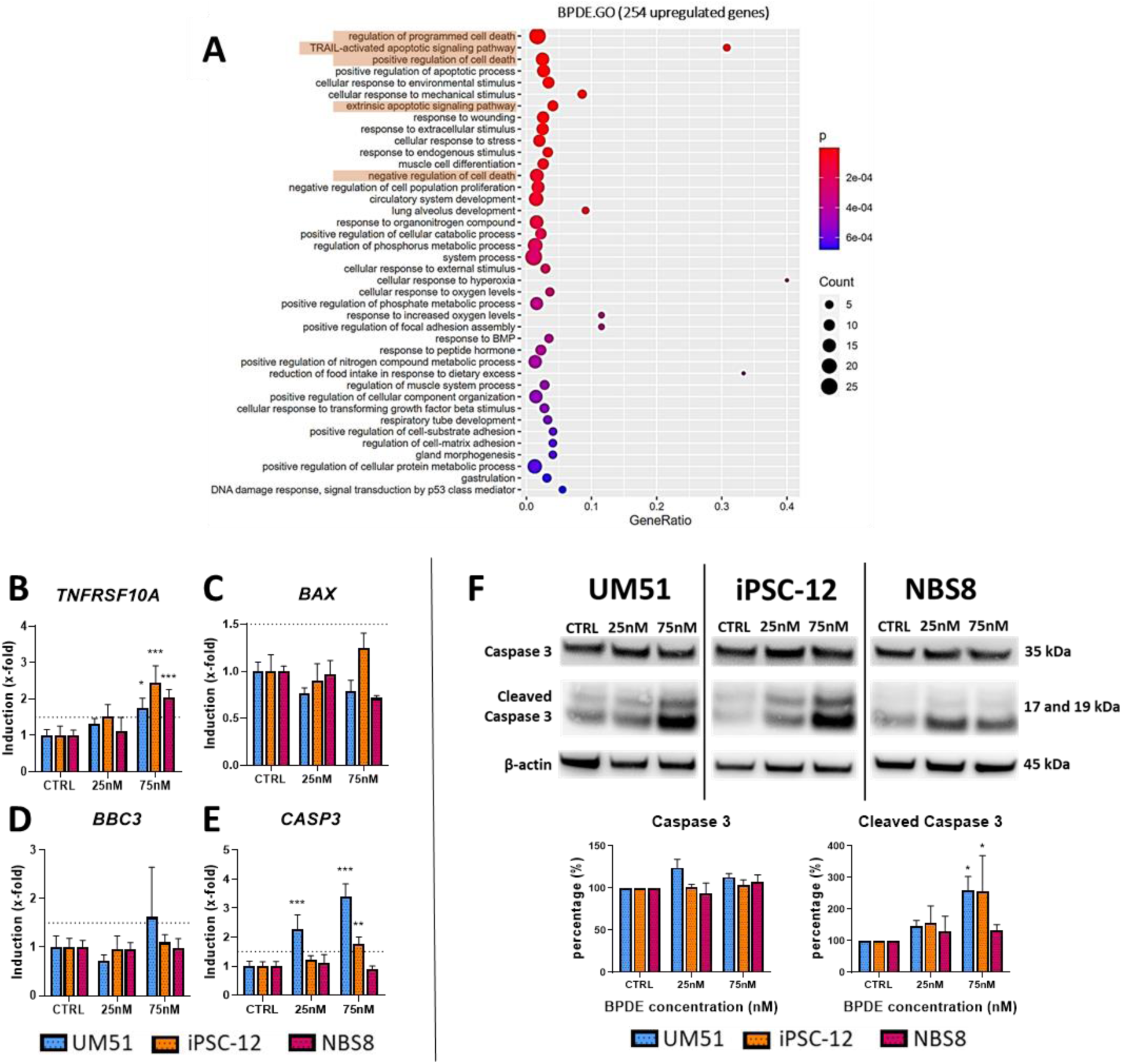
BPDE exposure differentially regulates apoptotic markers on WT and NBS8 hiPSCs. hiPSCs were treated with 25nM and 75nM of BPDE for 24h. RNA was extracted for microarray analysis and qRT-PCR. **(A)** Gene ontology analysis was performed for 254 genes upregulated in hiPSCs after BPDE treatment. Apoptosis related clusters are highlighted in orange. **(B, C, D, E)** qRT-PCR for *TNFRSF10A, BAX, BBC3*, and *CASP3*. Error bar depicts 95% confidence interval. N=3, *p<0.05, ** = p<0.01, *** = p<0.001. Dashed lines mark 1.5-fold. **(F)** Western blot for Caspase 3 and cleaved Caspase 3. β-actin was used as loading control. N=2, mean +/- standard deviation shown, *p<0.05.

Both the extrinsic and intrinsic apoptotic pathways depend on Caspases to induce cell death, therefore an executioner Caspase essential in both pathways-Caspase 3 (*CASP3*), was investigated. *CASP3* expression increased (mean > 1.7-fold) after BPDE treatment in both WT cell lines, but not in NBS8 (Figure 3B). Caspase 3 protein expression was unchanged in all three lines, but the expression of cleaved Caspase 3, the active form of Caspase 3, was enhanced particularly in the WT lines and far less in NBS8 (Figure 3F).

### 3.54 BPDE differentially regulates DNA damage response genes in WT and NBS8 hiPSCs

The enrichment analysis tool- Metascape, was used to compare the input gene list of the common 254 upregulated genes in all 3 cell lines after BPDE exposure. Two of the top non-redundant enrichment clusters were related to the p53 signaling pathway with 26 upregulated genes (Supplementary figure 3B). Among them were three known regulators of p53 signaling, *cJUN, MDM2* and protein Phosphatase, Mg^2+/^Mn^2+^ Dependent 1D (*PPM1D*), as well as other genes involved in p53-mediated DNA damage response such as *GADD45A*, Sestrin 1 (*SESN1*), *POLH*, BTG Anti-Proliferation Factor 2 (*BTG2*) and Stratifin (*SFN*).

As p53 is an important factor in the regulation of pluripotency and differentiation (35), we aimed to investigate the influence of BPDE exposure on p53 and its effectors in our model.

Interestingly, qRT-PCR revealed that *TP53* was significantly downregulated (mean < 0.5-fold) in both WT lines after treatment with 75nM BPDE (Figure 4A), whilst the expression p53 protein was increased in both healthy controls albeit only significantly in iPSC-12 hiPSCs (Figure 4B). Both gene and protein expression remained unchanged in NBS8 (Figure 4A and 4B). Two targets related to p53 regulation, *MDM2* and *cJUN*, were measured in an attempt to clarify the differences in the expression of *TP53* and p53 between WT and NBS8 cells after BPDE treatment. *MDM2* gene expression was downregulated (mean 0.56 -fold) exclusively in NBS8 (Figure 4A), while its protein levels were increased only in the WT cells (Figure 4B). A similar pattern was observed with cJUN. The mRNA levels (mean > 3.2 -fold) and protein expression were both increased in WT cells after treatment, but unchanged in NBS8 (Figure 4A and 4C). Since the activity of cJUN is stimulated by its phosphorylation at serines 63/73, the protein levels of serine 73 p-cJUN were also measured, this revealed increased expression in the iPSC-12 cells (Figure 4C). Next, we evaluated the mRNA expression of four upstream components of the p53 signaling cascade, *ATR*, *ATM,* and their respective downstream effectors, *CHEK1* and *CHEK2* (Supplementary figure 7). BPDE exposure did not affect the expression of *ATM*, *CHEK1* or *CHEK2* however, it upregulated the expression of ATR (mean 7.1 -fold) in the NBS8 cell line (Supplementary figure 7A). p53 is involved in NER, regulating the expression of targets such as XPC and DDB2, and it has also been implicated in the transcriptional regulation of trans-lesion synthesis DNA polymerases in reaction to DNA damage (173). After BPDE exposure, gene expression of the trans-lesion synthesis DNA polymerase *POLH* was upregulated (mean > 1.8-fold) in all three cell lines (Figure 4D). Interestingly, *XPC* and *DDB2* were upregulated exclusively in NBS8, and in both tested concentrations. *XPC* was upregulated by 1.5-fold in both concentrations, whilst the regulation of *DDB2* increased in a dose-dependent manner; 1.6-fold at 25nM and 3-fold at 75nM (Figure 4D).

**Figure 4:**
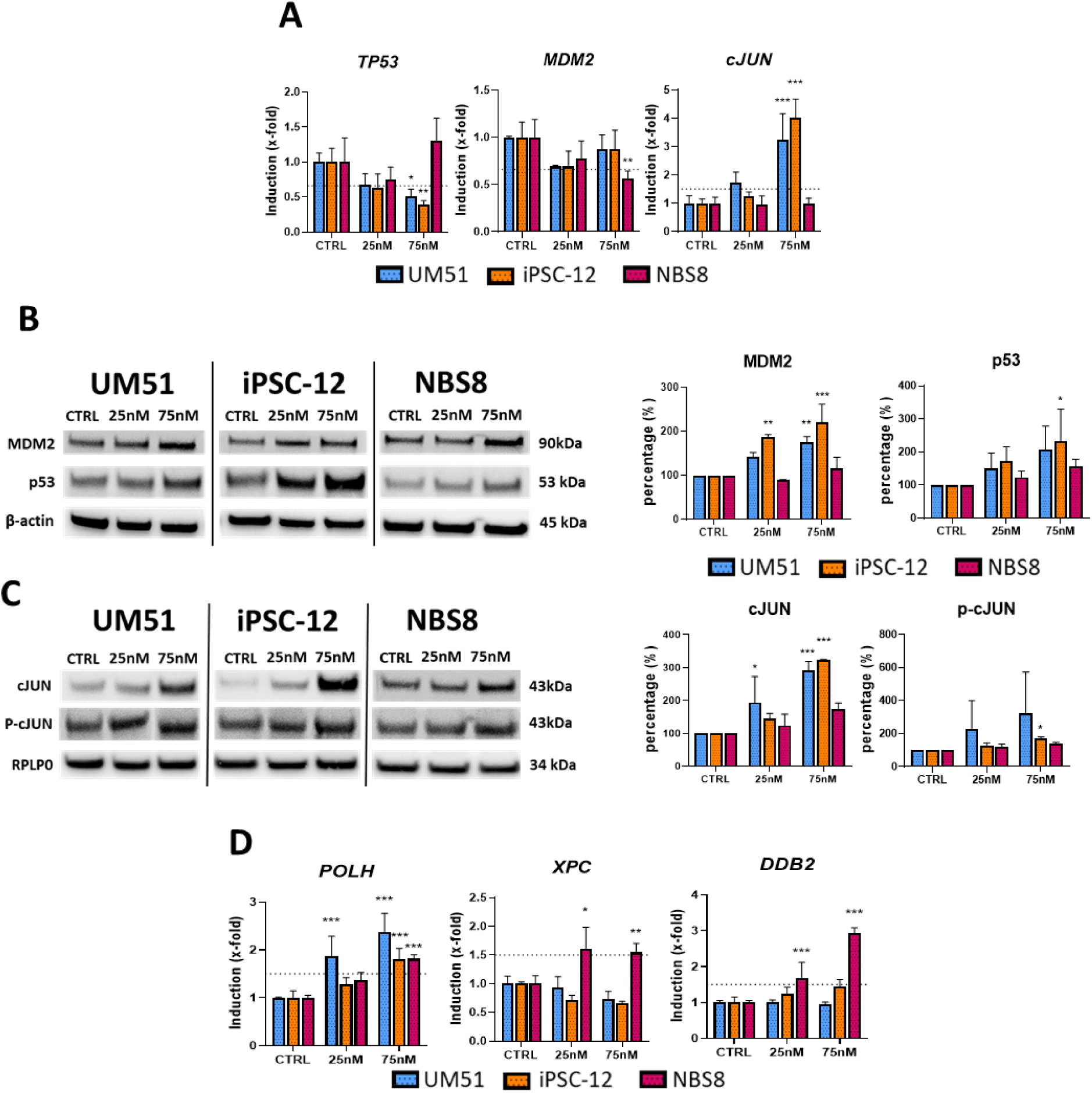
BPDE differentially regulates the DNA damage response in WT and NBS8 hiPSCs. hiPSCs were exposed to 25nM and 75nM of BPDE for 24h. **(A)** qRT-PCR for *TP53*, *MDM2* and *cJUN.* Error bar depicts 95% confidence interval. N=3, *p<0.05, ** = p<0.01. *** = p<0.001. Dashed lines mark 0.6 -fold (*TP53, MDM2*) and1.5-fold (*cJUN*). **(B and C)** Western blot for MDM2, p53, cJUN and p-cJUN. B-actin or RPLP0 were used as loading controls. N=2, mean +/- standard deviation shown, * = p<0.05, ** = p<0.01, *** = p<0.001. **(D)** qRT-PCR for *POLH*, *XPC* and *DDB2*. Error bar depicts 95% confidence interval. N=3, *p<0.05, ** = p<0.01, *** = p<0.001. Dashed lines mark 1.5-fold.

### 3.5. Responses to BPDE exposure is different in neural progenitor cells compared to their undifferentiated hiPSC parental lines

We next wanted to see, if BDPE has any influence on the early steps of neuro-development. As such, iPSCs were differentiated into NPCs and treated with 25nM or 75nM of BPDE for 24h. Immunostaining-based detection of expression of the NPC markers SOX2, SOX1 and Nestin, and the proliferation marker Ki67, was unchanged after BPDE treatment, and also for the neuron marker TUJ1 (Supplementary figure 8A). Likewise, gene expression levels of the NPC markers *SOX2*, *SOX1* and *PAX6* remained stable after genotoxic exposure (Supplementary figure 8B). Thus, BPDE treatment did not impair differentiation towards NPCs. However, they reacted less sensitive in terms of apoptosis and DNA damage marker expression than their undifferentiated counterparts.

The expression of *TNFRSF10A* and *CASP3* were upregulated in hiPSCs after BPDE treatment (Figure 3B and 3E), unlike in NPCs. The intrinsic apoptotic markers *BBC3* and *BAX* remained unchanged in both cell types (Figure 5A to 5D). The protein expression of Caspase 3, as well as that of its active form cleaved Caspase 3, remained mostly unchanged in NPCs after BPDE treatment, with only a slight upregulation (1.5-fold) of cleaved caspase 3 seen on iPSC-12 NPCs (Figure 5E). These results suggest that hiPSCs showed a lower resistance to BPDE cytotoxicity after 24h of exposure than their differentiated NPCs.

**Figure 5:**
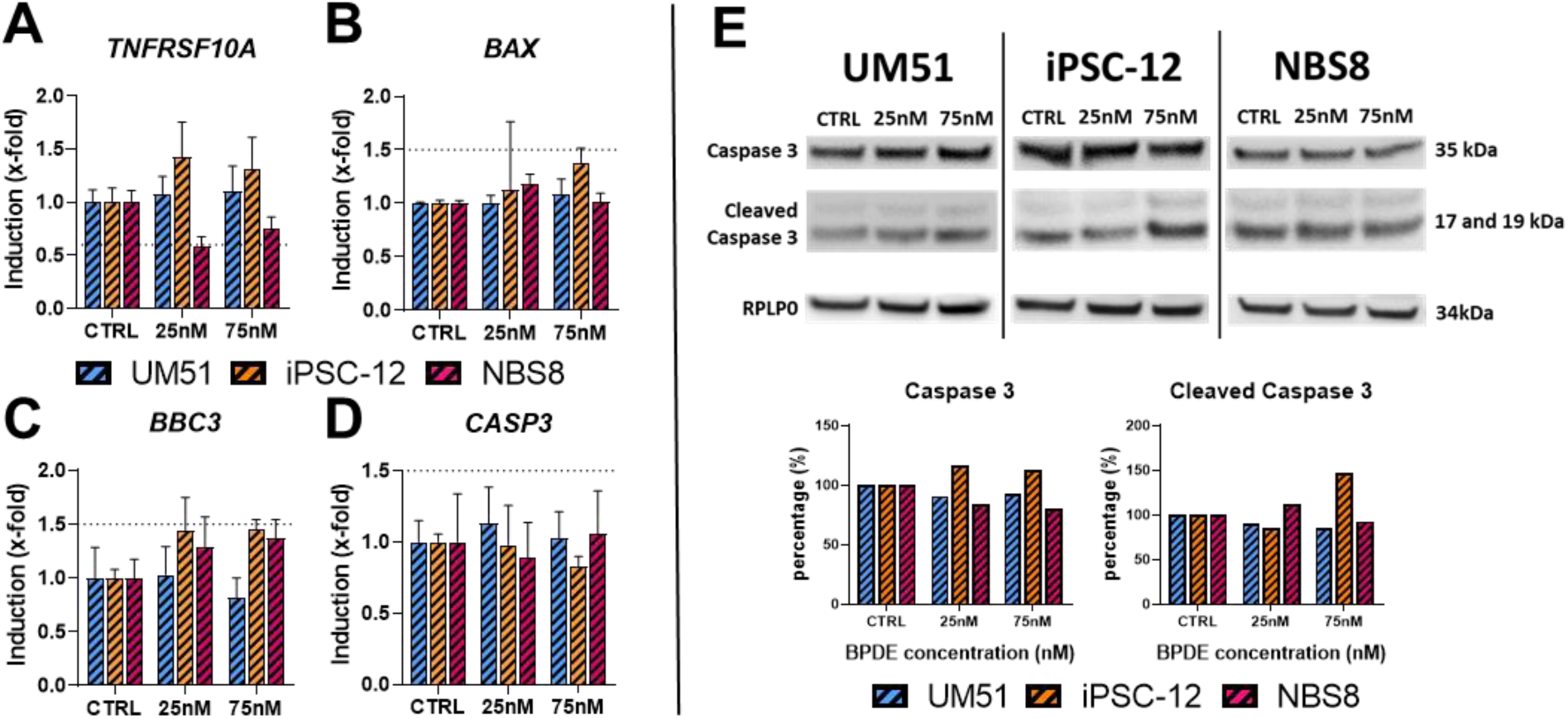
24h BPDE exposure does not influence expression of key apoptotic markers on NPCs. NPCs were treated with 25nM and 75nM of BPDE for 24h, then RNA was extracted for qRT-PCR and protein was extracted for western blot. qRT-PCR for **(A)** *TNFRSF10A*, **(B)** *BAX*, **(C)** *BBC3* and **(D)** *CASP3*. N=3. Error bar depicts 95% confidence interval. **(E)** Western blot and quantification for caspase 3 and cleaved caspase 3. N=1, RPLP0 was used for loading control.

Gene expression levels of the p53-inducible cell cycle regulators *CDKN1A* and *GADD45A* was unchanged in NPCs, with the exception of iPSC-12 NPCs, which showed a slight upregulation (1.8-fold) of both markers at 75nM (Figure 6A and 6B). With the intent to analyze the p53- mediated DNA damage response, the gene and protein expression of p53 was measured, as well as that of the p53 regulators MDM2 and cJUN. *TP53* and *MDM2* gene expression remained constant after BPDE treatment in NPCs in all lines, as did *cJUN* (Figure 6C to 6E). Investigation of the protein expression of MDM2 in NPCs after BPDE exposure revealed a similar pattern to that observed in hiPSCs, with the expression being enhanced in WT NPCs after treatment, but unchanged in NBS8 NPCs. Interestingly, p53 protein expression was enhanced only in iPSC-12 NPCs after genotoxic exposure, while the protein expression of cJUN was unchanged in all three NPC cell lines (Figure 6F).

**Figure 6:**
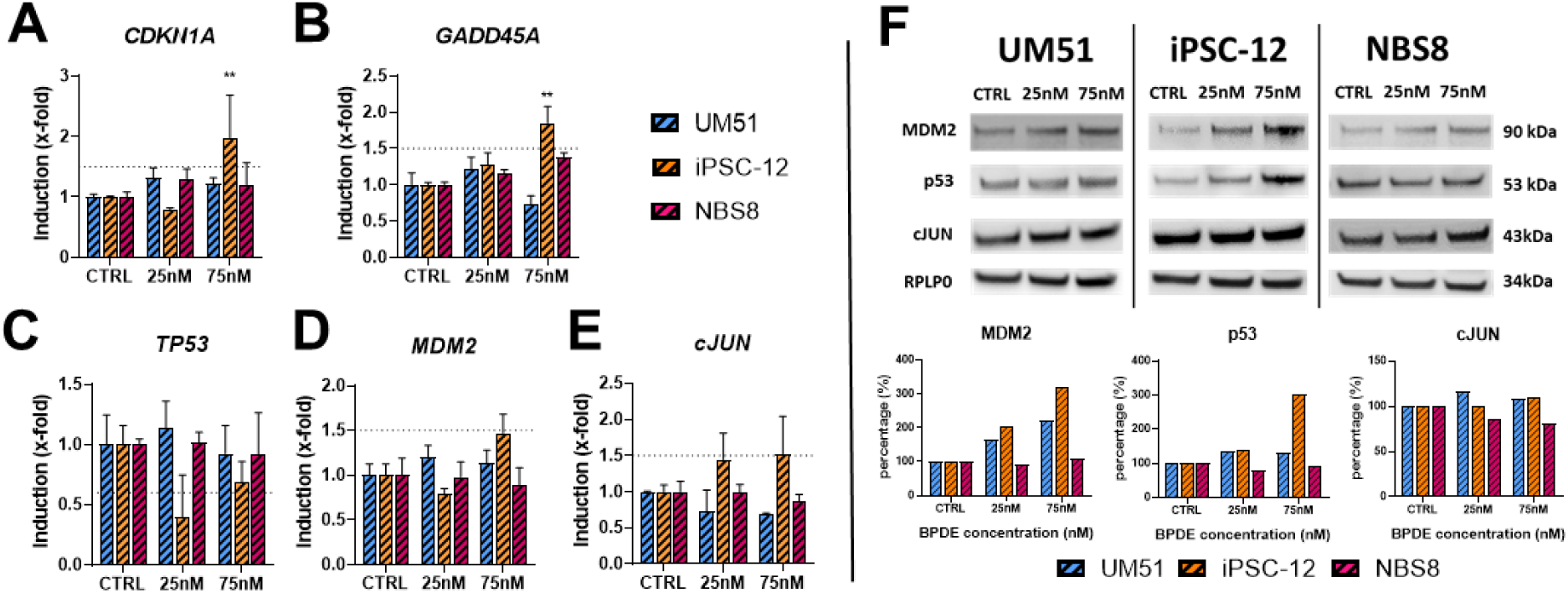
BPDE exposure of NPCs differentially regulates targets related to cell cycle control and p53 modulation when compared to hiPSCs. NPCs were treated with 25nM or 75nM of BPDE for 24h, then RNA was harvested for qRT-PCR and protein was harvested for western blot. qRT-PCR for (A) *CDKN1A*, (B) *GADD45A*, (C) *TP53* (D) *MDM2* and (E) *cJUN*. N=3, *p<0.05, **p<0.01. Error bar depict 95% confidence interval. Dashed lines mark 1.5- and 0.6-fold. **(F)** Western blot and quantification for MDM2, p53, and cJUN. N=1. RPLP0 was used as loading control.

While *ATM*, *CHEK1,* and *CHEK2* maintained a stable gene expression in both hiPSCs and NPCs after BPDE treatment, the upregulation of *ATR* which was noted in NBS8 hiPSCs after exposure to both 25nM and 75nM BPDE (1.6- fold and 7.1-fold respectively) was not observed in their differentiated NPC progeny (Supplementary figure 9A to 9D).

Lastly, gene expression of three p53-inducible genes related to bulky DNA-adduct repair, *POLH*, *XPC* and *DDB2*, was investigated. *XPC* and *DDB2* were upregulated in NBS8 hiPSCs in both concentrations tested (Figure 4D), but in NPCs *XPC* expression was unchanged after BPDE exposure (Supplementary figure 9F) and *DDB2* was only upregulated at 75nM (1.5-fold) and less than in their hiPSC counterparts (3-fold) (Supplementary figure 9E). In hiPSCs, the expression of *POLH* was upregulated in all three cell lines after BPDE treatment (Figure 5D), which was not seen in NPCs (Supplementary figure 9G).

These results are summarized in Table 1 and Table 2. An observation from the summary is that NBS8 cells, in particular NBS8 hiPSCs, present a distinct DNA damage response upon BPDE exposure than their WT counterparts. For example, NBS8 hiPSCs upregulate mRNA expression of *XPC* and DDB2, genes involved in NER, while showing a lower upregulation of targets involved in the apoptotic response such as cleaved Caspase 3. Secondly, the different cell types have distinct responses to BPDE exposure. hiPSCs have generally a more robust DNA damage response, with the regulation of several targets related to cell cycle checkpoint, apoptosis, and DNA damage repair, while NPCs had a more moderate response, with few regulated targets.

**Table 1:**
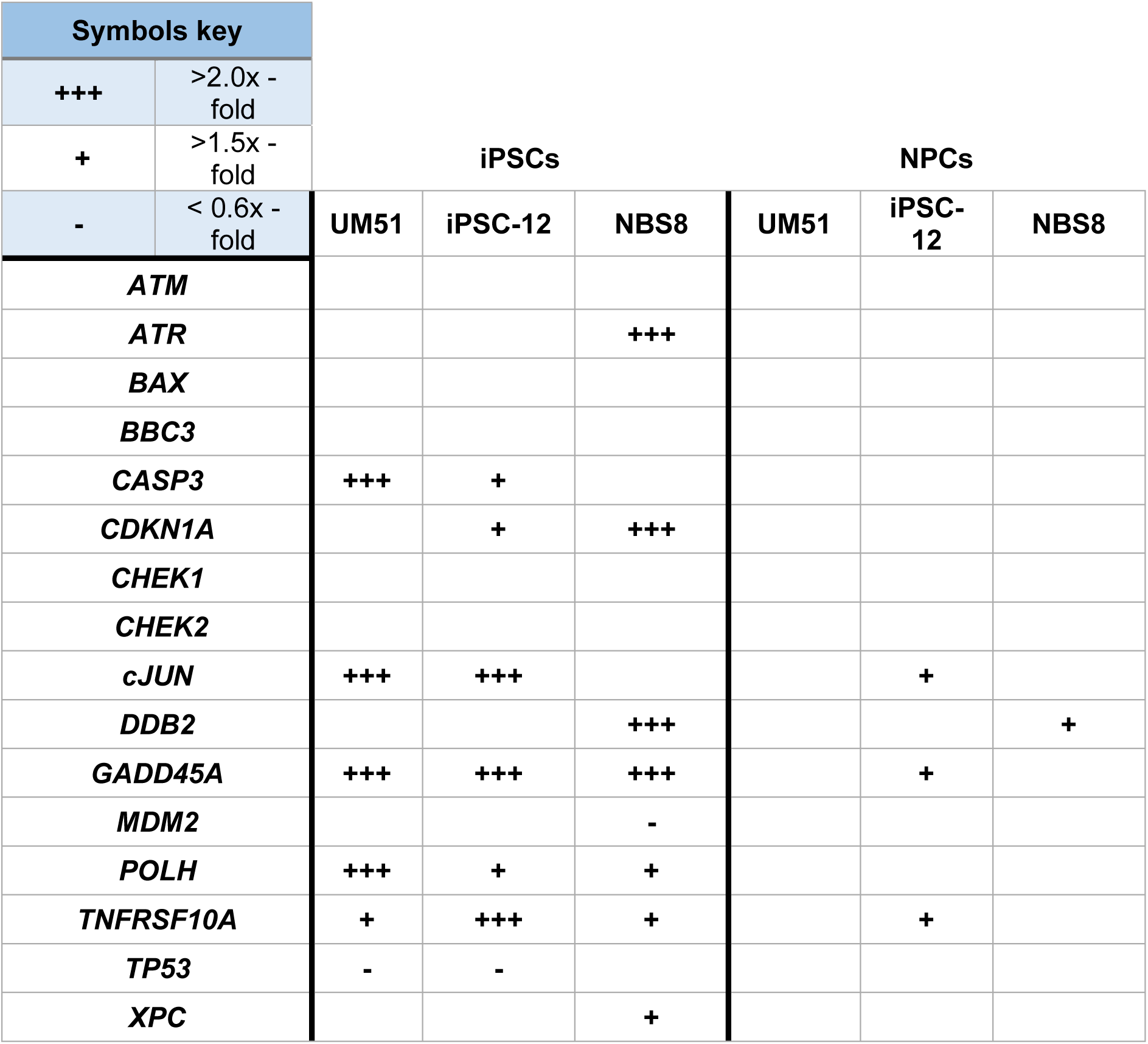
Summary table of results from the set of genes investigated through qRT-PCR after BPDE exposure. Cells were exposed to BPDE in the concentrations of 25nM and 75nM for 24h. Their RNA was extracted and used for qRT-PCR. The table summarizes the results obtained from the investigation of known BPDE targets as seen on the literature and those that were identified as BPDE targets through microarray analysis of BPDE exposed hiPSCs. Empty cells indicate no change in expression. One and two crosses indicate over 1.5x -fold or over 2x -fold increase in expression compared to control, respectively. The minus sign indicates a downregulation of at least less than 0.6x -fold.

**Table 2:** Summary table of results from the set of proteins investigated after BPDE exposure. Cells were exposed to BPDE in the concentrations of 25nM and 75nM for 24h. Then, their protein was extracted and used for western blot. The table summarizes the results obtained from the investigation of proteins related to the DNA damage response. Empty cells indicate no change in expression. One and two crosses indicate over 150% or over 200% increase in protein expression compared to control, respectively. Blocked cells indicates that the target was not investigated.

| Symbols key |  | iPSCs |  |  | NPCs |  |  |
| --- | --- | --- | --- | --- | --- | --- | --- |
| +++ | >200% increase | UM51 | iPSC-12 | NBS8 | UM51 | iPSC-12 | NBS8 |
| + | >150% increase |  |  |  |  |  |  |
| p53 |  | +++ | +++ |  |  | + |  |
| cJUN |  | +++ | +++ | + |  |  |  |
| P-cJUN |  | +++ | +++ |  |  |  |  |
| Caspase 3 |  |  |  |  |  |  |  |
| Cleaved Caspase 3 |  | +++ | +++ | + |  |  |  |
| MDM2 |  | +++ | +++ |  | +++ | +++ |  |

## 4. Discussion

To elucidate the effects of BPDE on undifferentiated hiPSCs at the transcriptional level, we performed for the first time a whole-genome transcriptome analysis of WT hiPSCs exposed to BPDE, which we compared to NBS patient-derived iPSCs that have a defect in the DNA repair mechanism.

Pluripotent stem cells have highly efficient DNA repair systems (36–38), but in case of failure to repair DNA lesions they will either rapidly undergo apoptosis (34) or suffer a p53-mediated loss of pluripotency by repressing the transcription factors *OCT4* and *NANOG*, and activating genes associated with differentiation (39,40). Our results show that 24h of BPDE exposure on hiPSCs affected neither the expression of *OCT4* and *NANOG*, nor that of targets associated with triggering differentiation such as *BMP4* or *GREM1* (Figure 1A and 1B). The nuclear expression of OCT4 also remained stable (Figure 1C). In line with data from Momcilovic et al, who studied the effects of γ-irradiation in hiPSCs (42), this suggests that our cells retained pluripotency for at least 24h after BPDE exposure.

BPDE treatment has been documented to induce cell cycle disruption in somatic cells, dependent on the treatment dose, the cell type, and the phase in the cell cycle that the cells were in at the start of treatment (41–43).

The GOs of hiPSC exposed to BPDE revealed an upregulated cluster related to DNA damage-induced cell cycle arrest. *GADD45A*, which is known to stimulate cell cycle arrest upon genotoxic exposure (44,45) was upregulated in all cell lines after treatment (Figure 2B). However, cell cycle analysis via flow cytometry for PI and EdU incorporation assays indicated that hiPSCs had no change in their cell cycle distribution after BPDE exposure (Figure 2C and 2D). This could be related to the timepoint of the analysis, since it has been shown that hiPSCs exposed to γ-irradiation show cell cycle arrest at the G2/M phase in the first 9h after exposure, but after 24h their cell cycle distribution returns to the same pattern as non-irradiated controls (37). Interestingly, *CDKN1A*, a p53-associated cell cycle regulator (46), was up-regulated only in the NBS8 cell line after BPDE exposure (Figure 2A), lending support to reports of perturbed cell cycle arrest signaling after genotoxic exposure in NBN-impaired cells (47–49).

Bioinformatic analysis of hiPSC exposed to BPDE revealed upregulated clusters related to the p53 signalling pathway (Supplementary Figure 3B). Investigation of the gene and protein expression of p53, MDM2 and cJUN, as well as Ser73 phospho-cJUN, revealed a general pattern of upregulation of these targets in WT hiPSC and little or no regulation in NBS8 cells (Figure 4).

Although *TP53* gene expression was downregulated in WT hiPSCs after BPDE exposure, p53 protein expression was increased in these cells, while NBS8 hiPSCs showed no *TP53* or p53 regulation (Figure 4A). Interestingly, dysregulation of p53 function upon DNA damage is a defining aspect of NBS. It is known that NBS fibroblasts exposed to ionizing radiation have a delayed and reduced p53-mediated response to DNA damage (51). We made a similar observation for NBS fibroblasts and NBS iPSC-derived cerebral organoids exposed to bleomycin in earlier studies (50,51). This effect does not seem to be related to MDM2 mediated p53 degradation, as MDM2 protein expression was significantly increased only in WT hiPSCs after BPDE treatment but not in NBS8 hiPSC (Figure 4A and 4B).

The upregulation of cJUN and Ser73 p-cJUN in BPDE treated WT hiPSCs (Figure 4A and 4C) could help explain the downregulation of *TP53* and the low or no upregulation of *CDKN1A* mRNA levels seen in the cells and hint at a protective effect being elicited. cJUN has a variety of roles during the DNA damage response, repressing p53 mRNA expression to allow for cell cycle progression and proliferation (52), and playing a role in genotoxic resistance and activation of DNA damage repair (53–55). It is interesting to note that the phosphorylation of cJUN at Ser63/73 seems to be essential for its protective role against DNA damaging agents (54,55). However, cJUN has also been linked to cellular stress induced apoptosis, which seems to occur through the sustained upregulation of cJUN levels and, at least in some cell types, enhanced extrinsic apoptotic signalling (56,57). Thus, a possible pro-apoptotic effect through the extrinsic apoptotic pathway cannot be discarded.

Indeed, GO analysis of genes which were upregulated after BPDE exposure on hiPSCs revealed several clusters associated with positive and negative regulation of apoptosis (Figure 3A). Curiously, while reports of BPDE exposure on somatic cells reported the upregulation of the intrinsic apoptotic pathway, and genes such as *BAX* and *BBC3* (58–60), hiPSCs showed a distinct upregulation of the extrinsic apoptotic pathway, confirmed through qRT-PCR (Figure 3A to 3C). It was also observed that, although the gene expression of *TNFRSF10A* was upregulated in all three cell lines, the expression of the executioner *CASP3* was only upregulated in WT cells (Figure 3D), and the same was observed for the protein expression of cleaved Caspase 3, the active form of the Caspase 3 protein (Figure 3E). NBN-impaired cells have been reported as having deficient apoptosis regulation (47,50). This translates into a delayed genotoxic exposure response that leads to deficient activation of the apoptotic pathway after DNA damage (47–49).

In line with this, our data confirm previously reported delays in the activation of the DNA repair machinery in NBN-deficient hiPSCs after BPDE treatment (47–49), as we could only observe up-regulation of X*PC* and *DDB2* in these cells but not in the WT cells (Figure 4B and 4C). *POLH* expression, however, was upregulated in all three hiPSC lines (Figure 4A). *POLH* is involved in the error-prone bypass of BPDE-adducts (61,62) and induction of *POLH* by BPDE has been implicated in enhanced cell survival, but at the expense of a higher number of genomic mutations (41,58). This adaptative reaction is particularly worth of further study since mutations that occur during early embryogenesis can contribute to cancer development later in life (63).

## 5. Conclusion

To the best of our knowledge, this is the first time the effects of BPDE exposure have been investigated on an hiPSCs model and the first comparison made between the effects of BPDE on pluripotent stem cells and differentiated NPCs. This mimics pre-gastrulation and differentiation towards the ectoderm lineage of early human development.

We showed that hiPSC and NPCs harbouring an NBS mutation reacted differently to BPDE treatment compared to WT cells, showing less apoptotic response and no increase in the expression of p53 and MDM2. Our data also emphasizes the differences in the DNA damage response between hiPSCS and NPCs, with the former presenting a robust response compared to NPCs, enhancing the mRNA and/or protein expression of several targets related to DNA damage response, apoptosis and cell cycle checkpoints. Furthermore, our model conforms with the 3Rs principle.

## Supplementary Materials

**Supplementary figure 1: Dose-response curve of BPDE-treated hiPSCs.**

**Supplementary figure 2: Venn Diagrams of differentially regulated genes extracted from microarray analysis.**

**Supplementary figure 3: Metascape analysis of differentially regulated genes extracted from microarray analysis.**

**Supplementary figure 4: GO and KEGG pathways of differentially regulated genes extracted from microarray analysis.**

**Supplementary figure 5: In control conditions, NBS8 hiPSCs have enhanced cancer related KEGG pathways compared to WT hiPSCs.**

**Supplementary figure 6: BPDE treatment enhances cancer-related GO clusters and KEGG pathways in NBS8 hiPSCs.**

**Supplementary figure 7: BPDE differentially regulates the gene expression of DNA damage response gene *ATR* between WT and mutant hiPSCs.**

**Supplementary figure 8: 24h BPDE exposure on NPCs does not affect expression of key NPC markers.**

**Supplementary figure 9: BPDE exposure differentially regulates gene targets related to DNA damage response and repair in NPCs when compared to hiPSCs.**

**Supplementary table 1: Cell lines used in this work.**

**Supplementary table 2: Antibodies and dilutions used on immunocytochemistry (ICC) and Western blot.**

**Supplementary table 3: Primer sequences used in this work.**

## Institutional Review Board Statement

The study was conducted according to the guidelines by the Ethics Committee of the medical faculty of Heinrich-Heine University, Germany (protocol code: 5704).

## Data availability statement

All microarray data will be available at NCBI GEO server once the manuscript is accepted.

