## Supplementary Data for "Investigating BPDE-induced embryonic toxicity employing hiPSC-based models"

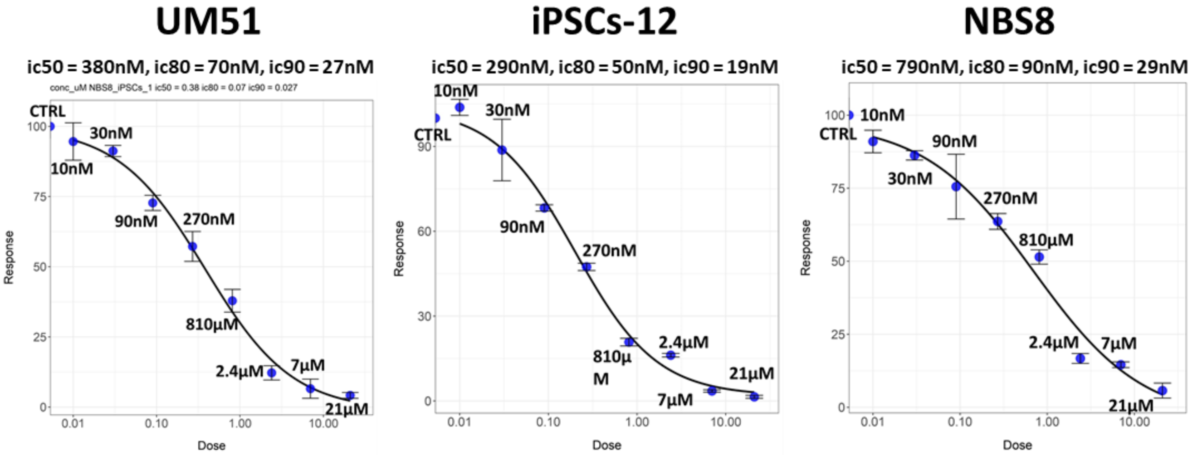

**Supplementary figure 1: Dose-response curve of BPDE-treated hiPSCs.** hiPSCs were treated with increasing concentrations of BPDE for 24h and resazurin assay was performed to measure cell viability. Dot plot graph represents the dose-response curve, with each dot representing the mean of three biological triplicates. +/- standard deviation is shown.

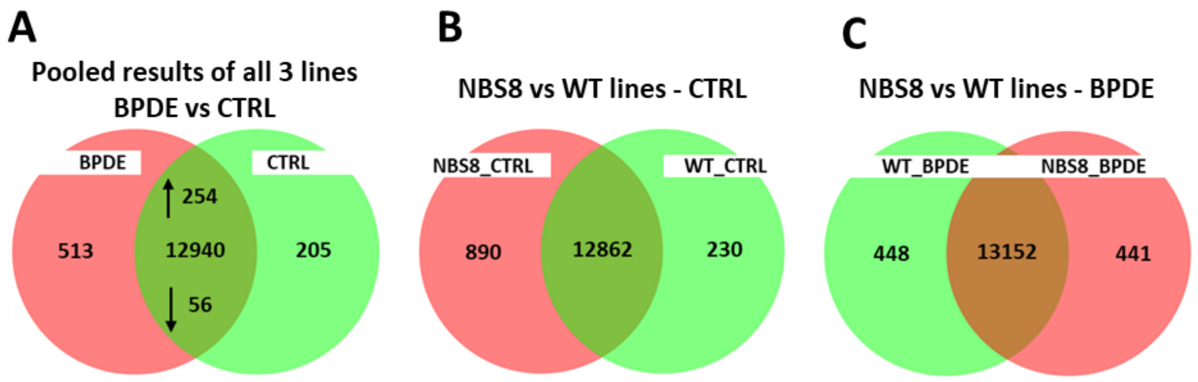

**Supplementary figure 2: Venn Diagrams of differentially regulated genes extracted from microarray analysis.** WT and NBS8 hiPSCs were treated with 75nM BPDE for 24h, then RNA was harvested and subjected to Affymetrix analysis. The expression profiles were extracted and organized regarding the regulation of **(A)** BPDE treatment on hiPSCs as well as the differences between the WT and mutant lines, both in **(B)** control and **(C)** treated conditions**.**

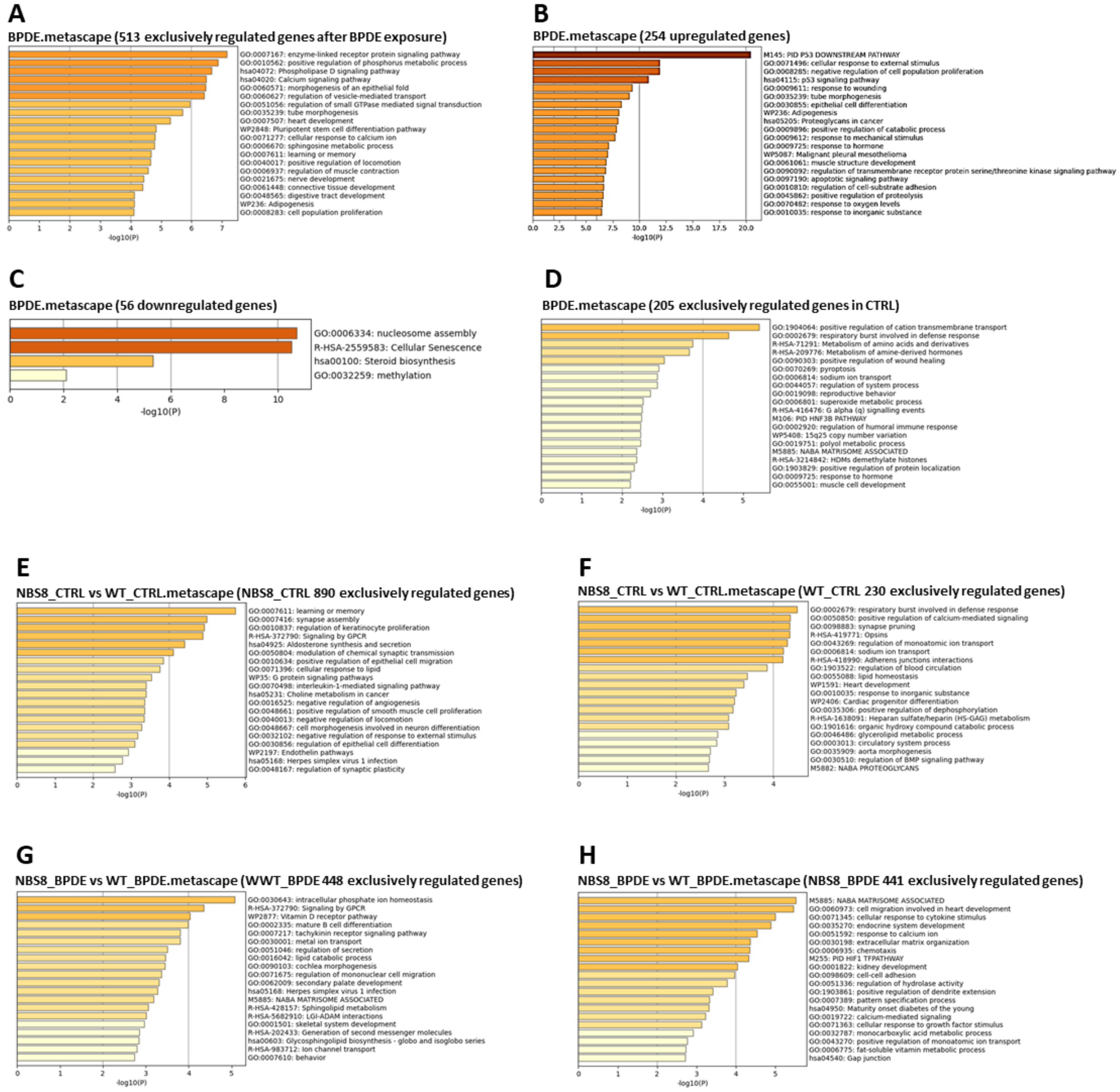

**Supplementary figure 3: Metascape analysis of differentially regulated genes extracted from microarray analysis.** WT and NBS8 hiPSCs were treated with 75nM BPDE for 24h, then harvested and subjected to Affymetrix analysis. The expression profiles were extracted, and the identified differentially regulated genes were investigated using metascape: **(A to D)** Differentially regulated genes of three hiPSC lines subjected to BPDE exposure. **(E and F)** Exclusively regulated genes of WT and NBS8 lines in control conditions. **(G and H)** Exclusively regulated genes of WT and NBS8 lines after BPDE exposure.

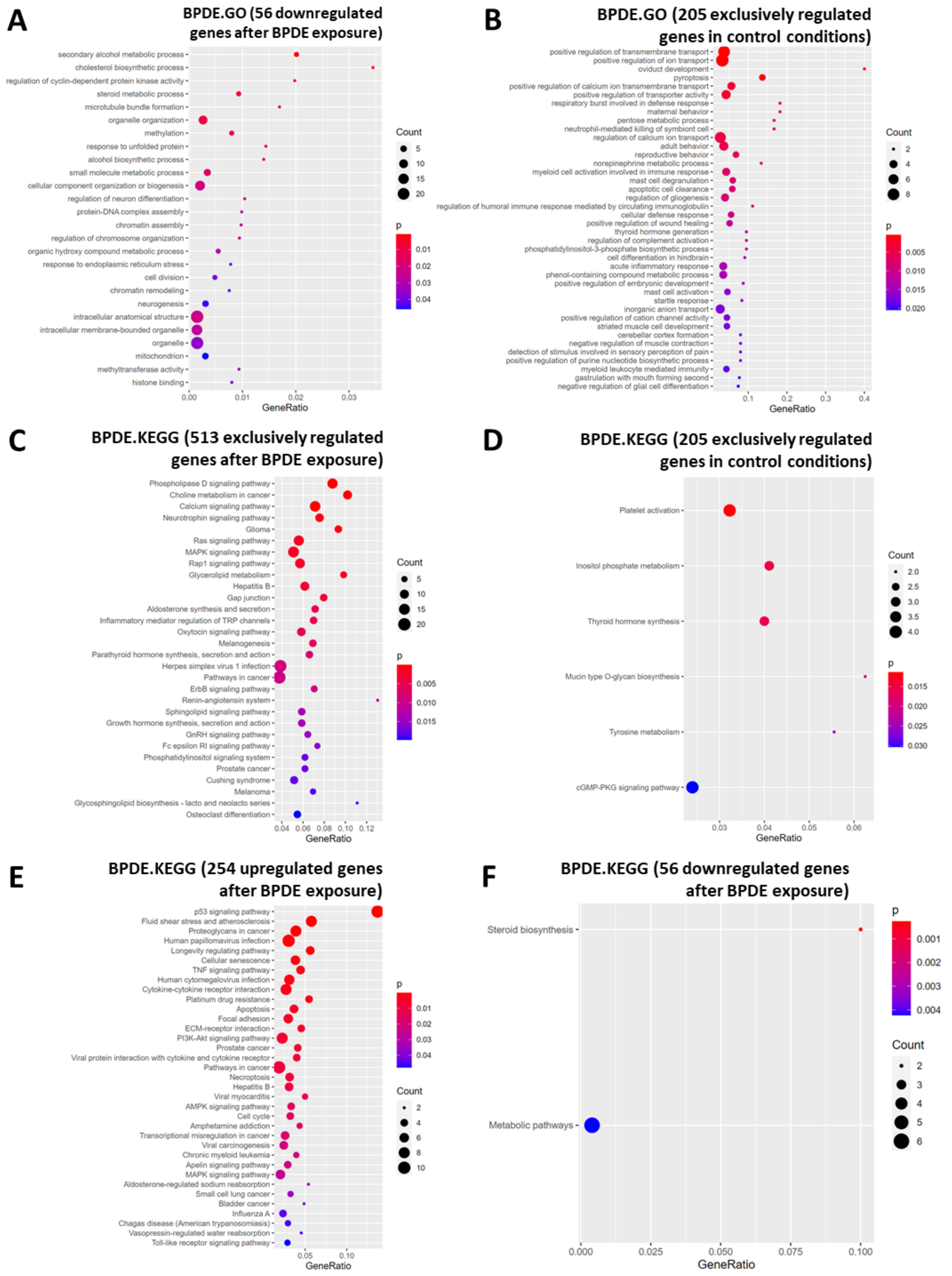

**Supplementary figure 4: GO and KEGG pathways of differentially regulated genes extracted from microarray analysis.** WT and NBS8 hiPSCs were treated with 75nM BPDE for 24h, then harvested and subjected to Affymetrix analysis. The expression profiles were extracted, the results of all three iPSCs pooled together and the identified differentially regulated genes were investigated using GO and KEGG pathways. **(A)** GO from 56 genes downregulated in hiPSCs after BPDE exposure. **(B)** GO from 205 genes exclusively regulated in hiPSCs in control conditions. **(C)** KEGG pathways from 513 genes exclusively regulated in hiPSCs after BPDE exposure. **(D)** KEGG pathways from 205 genes exclusively regulated in hiPSCs in control conditions. **(E)** KEGG pathways from 254 genes upregulated in hiPSCs after BPDE exposure. **(F)** KEGG pathways from 56 genes downregulated in hiPSCs after BPDE exposure.

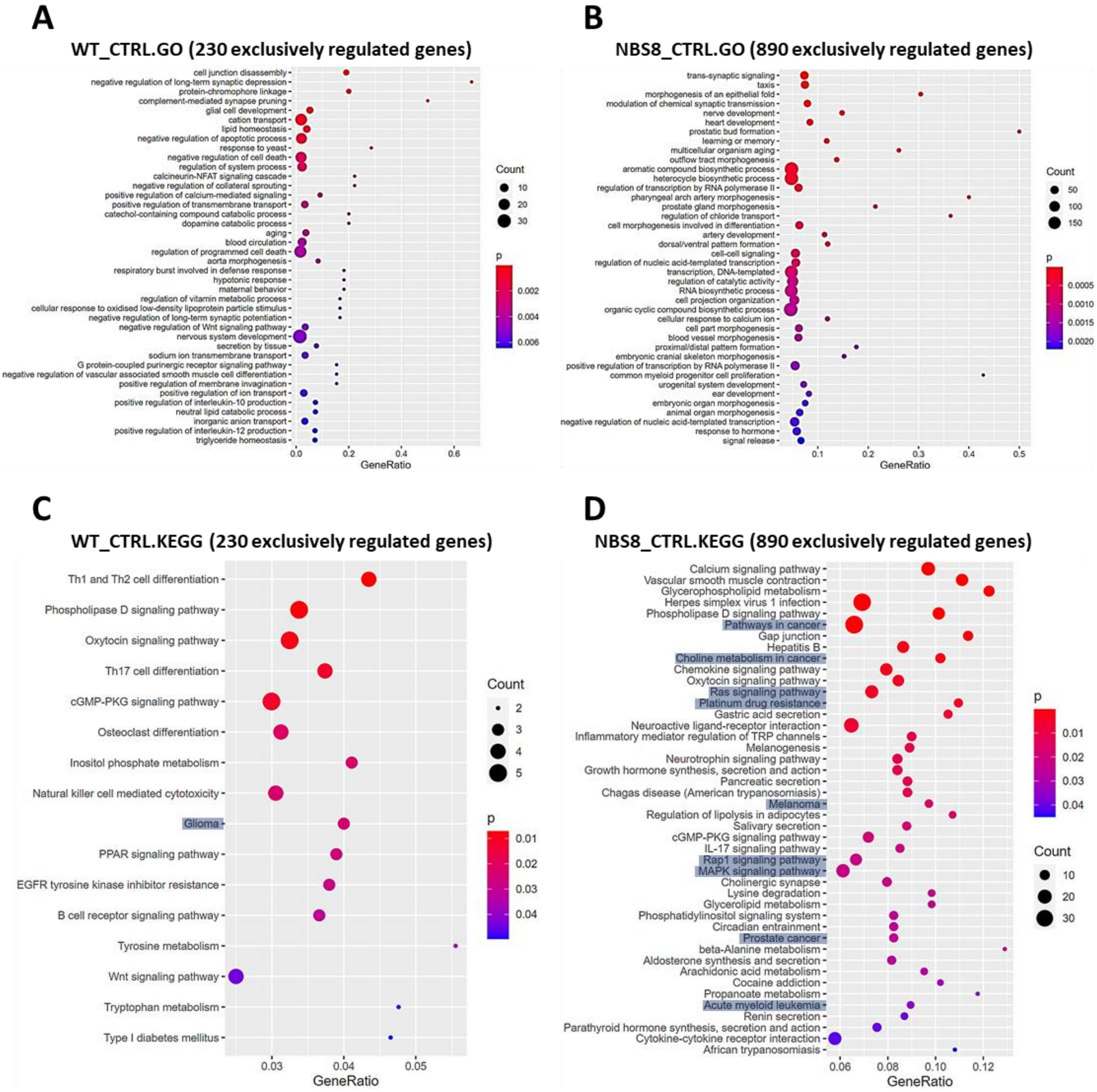

**Supplementary figure 5: In control conditions, NBS8 hiPSCs have enhanced cancer related KEGG pathways compared to WT hiPSCs.** hiPSCs were treated with 75nM of BPDE for 24h, then had their RNA harvested for affimetrix microarray. **(A)** GO from 230 genes exclusively regulated in WT hiPSCs in control conditions. **(B)** GO from 890 genes exclusively regulated in NBS8 hiPSCs in control conditions. **(C)** KEGG pathways from 230 genes exclusively regulated in WT hiPSCs in control conditions. **(D)** KEGG pathways from 890 genes exclusively regulated in NBS8 hiPSCs in control conditions.

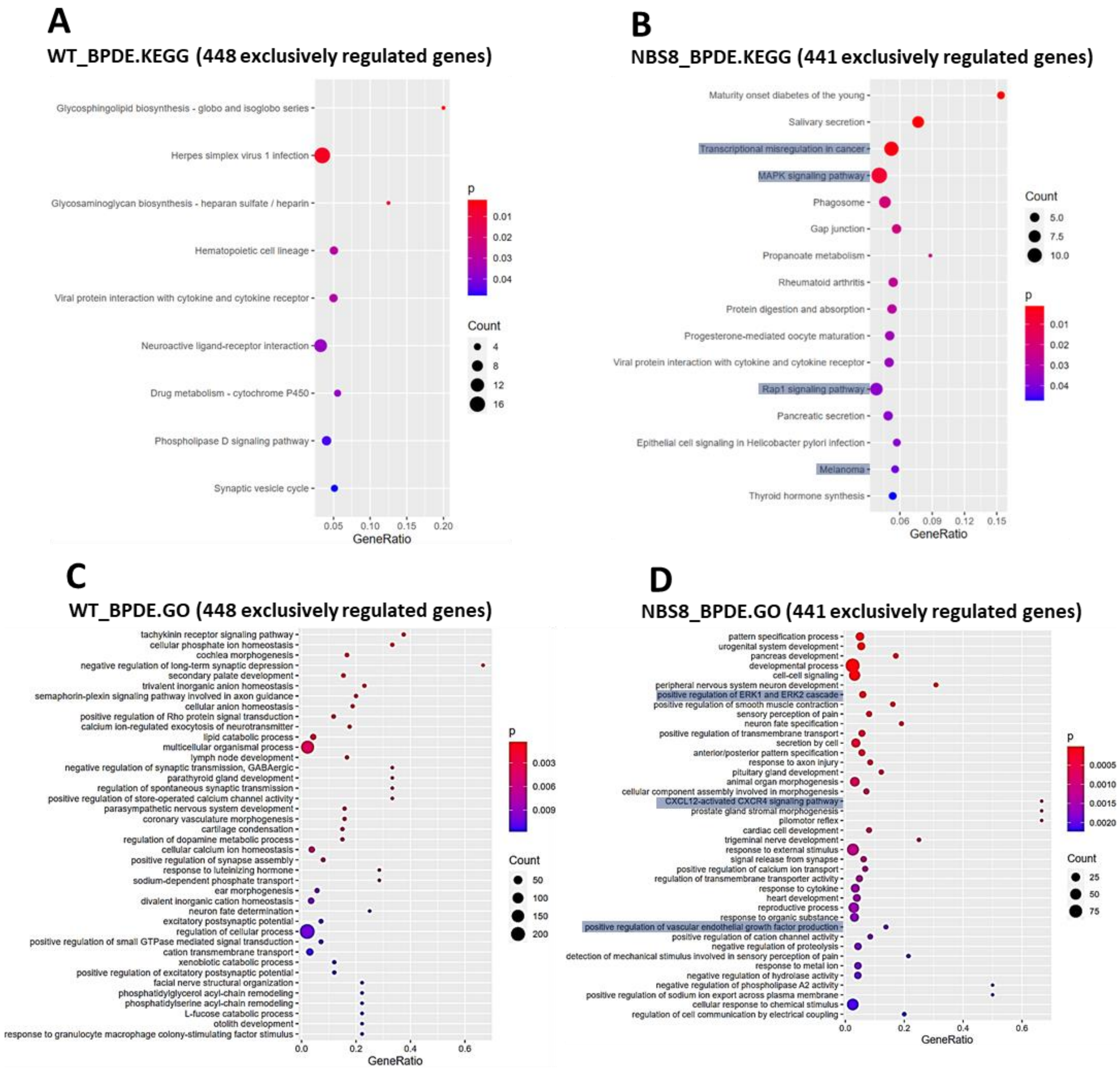

**Supplementary figure 6: BPDE treatment enhances cancer-related GO clusters and KEGG pathways in NBS8 hiPSCs.** hiPSCs were treated with 75nM of BPDE for 24h, then had their RNA harvested for affimetrix microarray. **A)** KEGG pathways from 448 genes exclusively regulated in WT hiPSCs after BPDE treatment. **(B)** KEGG pathways from 441 genes exclusively regulated in NBS8 hiPSCs after BPDE treatment. **(C)** GO from 448 genes exclusively regulated in WT hiPSCs after BPDE treatment. **(D)** GO from 441 genes exclusively regulated in NBS8 hiPSCs after BPDE treatment.

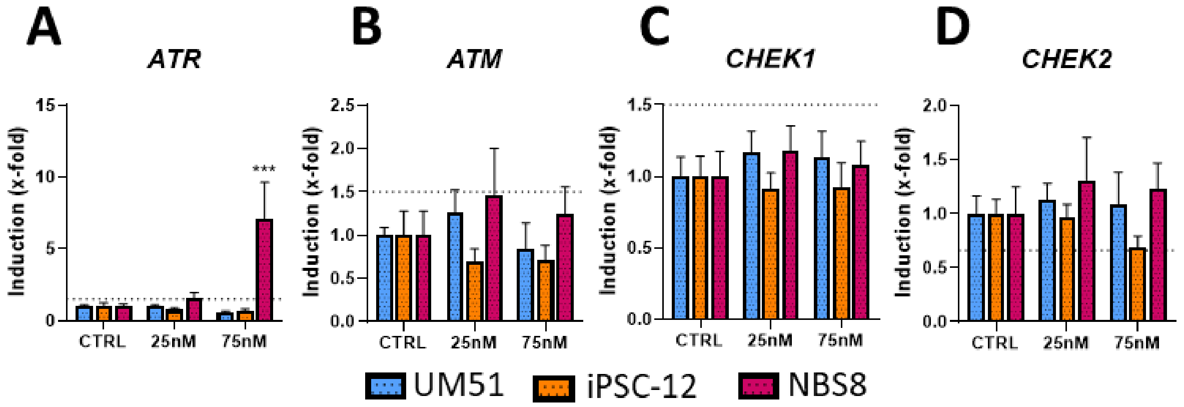

**Supplementary figure 7: BPDE differentially regulates the gene expression of DNA damage response gene *ATR* between WT and mutant hiPSCs.** hiPSCs were exposed to 25nM and 75nM of BPDE for 24h, then RNA was extracted for qRT-PCR for **(A)** *ATR*, **(B)** *ATM*, **(C)** *CHEK1* and **(D)** *CHEK2*. Error bar depicts 95% confidence interval. N=3, *p<0.05, ** = p<0.01. *** = p<0.001. Dashed lines mark 0.6 -fold and1.5-fold.

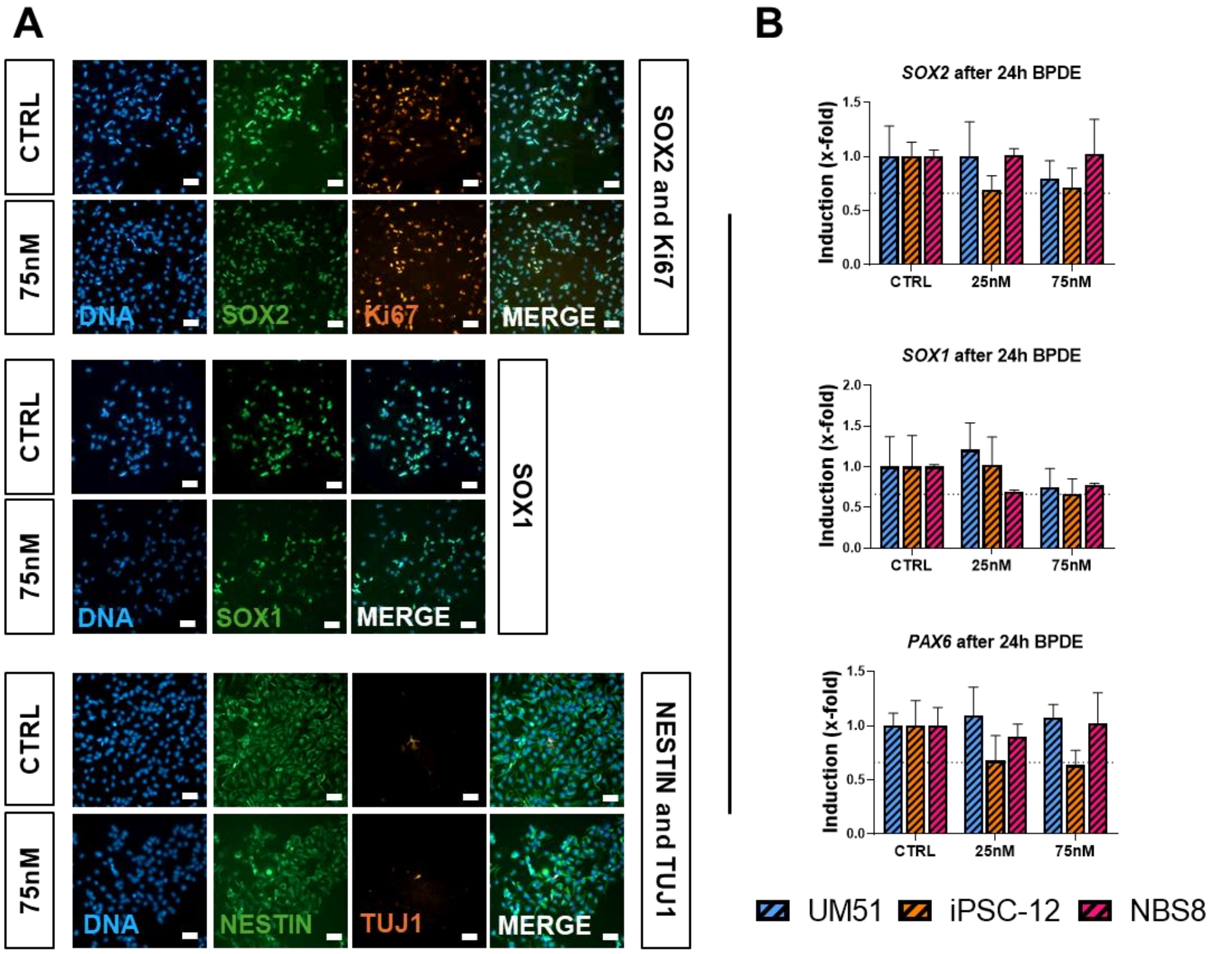

**Supplementary figure 8: 24h BPDE exposure on NPCs does not affect expression of key NPC markers.** NPCs were exposed to BPDE at 25nM and 75nM concentrations for 24h, then RNA was harvested for qRT-PCR or cells were fixed for immunocytochemistry. **(A)** Immunocytochemistry for NPC markers SOX2, Ki67, SOX1, Nestin and the neuron marker TUJ1. Scale bar 50µm. **(B)** qRT-PCR for NPC markers *SOX2*, *SOX1* and *PAX6*. N=3. Error bar depicts 95% confidence interval. Dashed lines mark 0.6 -fold.

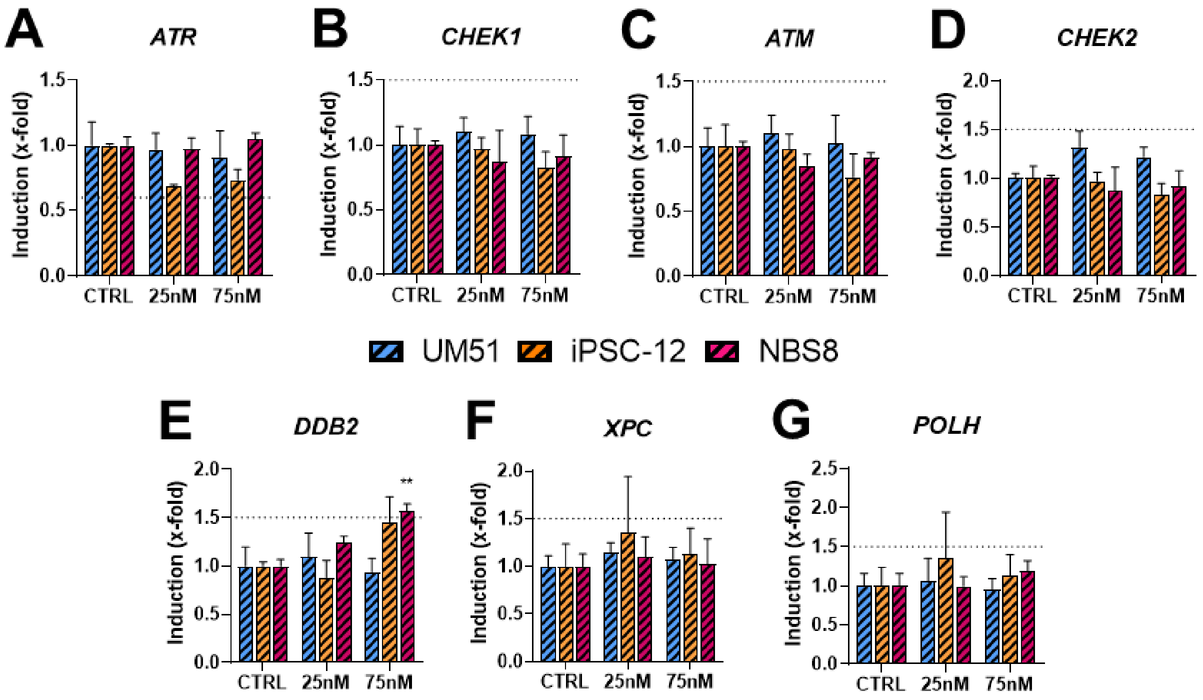

**Supplementary figure 9: BPDE exposure differentially regulates gene targets related to DNA damage response and repair in NPCs when compared to hiPSCs.** NPCs were treated with 25nM and 75nM of BPDE for 24h and RNA was harvested for qRT-PCR for **(A)** *ATR*, **(B)** *CHEK1,* **(C)** *ATM*, **(D)** *CHEK2*, **(E)** *DDB2,* **(F)** *XPC*, **(G)** and *POLH*. N=3, *p<0,05, **p<0,01 ***p<0,001. Error bar depicts 95% confidence interval. Dashed lines mark 1.5-fold.

**Supplementary table 1: Cell lines used in this work.** M: male, F: female.

| Cell line | Gender | Genotype | Age of Donor | Derived from | Reprogramming method | Source |
|---|---|---|---|---|---|---|
| ISRM-UM51 | M | - | 51 years old | SIX2-positive renal cells | Episomal reprogramming | (1) |
| iPSC-12 | F | - | Neonatal | Dermal fibroblast | Retroviral transduction | Cell applications Inc. |
| NBS8 | F | Heterozygous NBN 657del5 | 7 years old | Dermal fibroblasts | Retroviral transduction | (2) |

**Supplementary table 2: Antibodies and dilutions used on immunocytochemistry (ICC) and Western blot.** For ICC, all antibodies were diluted in 3%BSA/DPBS unless stated otherwise. For western blot, all antibodies were diluted in 5% BSA/TBS-T unless stated otherwise. CST: Cell signaling Technologies. R&D: R&D Systems. SySy: Synaptic Systems. gp: guinea pig; gt: goat; ms: mouse; rb: rabbit. MW: expected molecular weight of protein band.

| Antibody | Brand | ICC | Western Blot | ID number |
|---|---|---|---|---|
| **ms Tuj1** | CST | 1:750 | - | TU-20 |
| **ms Ki67** | CST | 1:200 | - | 9449S |
| **ms p53** | CST | | 1:1000 in 5% milk/TBS-T. (MW: 53kDa) | mAb2524 |
| **rb Nestin** | SigmaAldrich | 1:1000 | | N5413 |
| **ms b-Actin** | CST | | 1:5000 in 5% milk/TBS-T (MW:45kDa) | 3700S |
| **rb cJUN** | CST | | 1:1000. (MW: 43 kDa) | 9165S |
| **rb Cleaved Caspase 3** | CST | | 1:1000. (MW: 17-19 kDa) | 9664S |
| **rb anti p-cJUN** | CST | | 1:1000. (MW: 43 kDa) | 3270S |
| **rb anti SOX2** | CST | 1:400 | - | 3579S |
| **rb CASPASE 3** | CST | | 1:1000. (MW: 35 kDa) | 9662S |
| **rb MDM2** | CST | | 1:1000 (90 kDa) | 86934S |

| | | | | |
|---|---|---|---|---|
| **rb RPLP0** | Proteintech | | 1:1000 (34kDa) | 11290-2-AP |

**Supplementary table 3: Primer sequences used in this work.** All primers were produced by Eurofins Scientific.

| | | |
|---|---|---|
| ***ATM*** | TTTTCAACCAGTTTTCCGTTACTT C | ACACTGCGCGTATAAGCCAATC |
| ***ATR*** | CTGCCACTCAGCTTACCACT | AAGCTGTGCTGGGCTACATT |
| ***BBC3*** | TCCTGGGTCCCTGGCCAAGAAG | GTGTCACCCCTGCAGCTGGAAC |
| ***CASPASE 3*** | TCATTATTCAGGCCTGCCGTGGT A | TGGATGAACCAGGAGCCATCCTT T |
| ***CDKN1A*** | GATGTCCGTCAGAACCCATGCG | GTCGAAGTTCCATCGCTCACGG |
| ***CHEK1*** | CAAGAAAGGGGCAAAAAGG | TGTATGAGGGGCTGGTATCC |
| ***CHEK2*** | TTCAGCAAGAGAGGCAGACC | GCGTTTATTCCCCACCACTT |
| ***DDB2*** | GCCATCTGTCCAGCAGGGGC | GGGGTGAGTTGGGTGCCACG |
| ***cJUN*** | ACCTTGAAAGCTCAGAACTCGG | TTAGCATGAGTTGGCACCCAC |
| ***GADD45A*** | CAGGCGTTTTGCTGCGAGAACG | TGTGGATTCGTCACCAGCACGC |
| ***MDM2*** | AAACTGGGGAGTCTTGAGGG | TGCACATTTGCCTGCTCCTC |
| ***POLH*** | GCCCACAACAGCCAAAGCATGC | GGGGTTTGAAGAGTGGGGCTGC |
| ***RPLP0*** | TCGACAATGGCAGCATCTAC | ATCCGTCTCCACAGACAAGG |
| ***SOX1*** | TTGGCATCTAGGTCTTGGCTCA | CGGGCGCACTAACTCAGCTT |
| ***SOX2*** | GTATCAGGAGTTGTCAAGGCAGA G | TCCTAGTCTTAAAGAGGCAGCAA AC |
| ***TNFRSF10 A*** | CACACCCTGCTGGATGCCTTGG | CAAGGACACGGCAGAGCCTGTG |
| ***TP53*** | CAGGGCAGCTACGGTTTC C | CAGTTGGCAAAACATCTTGTTGAG |
| ***TUJ1*** | ATGAACACCTTCAGCGTCGT | CATCCGTGTTTTCCACCAGC |
| ***XPC*** | GACCTCAAGAAGGCACACCA | TGGCTTCACAGGCAGAAGAG |
